# Taxonomic bias in AMP prediction of invertebrate peptides

**DOI:** 10.1101/2021.03.04.433925

**Authors:** Zoltán Rádai, Johanna Kiss, Nikoletta A. Nagy

## Abstract

Invertebrate antimicrobial peptides (AMPs) are at the forefront in the search for agents of therapeutic utility against multi-resistant microbial pathogens, and in recent years substantial advances took place in the *in silico* prediction of antimicrobial function of amino acid sequences. A yet neglected aspect is taxonomic bias in the performance of these tools. Owing to differences in the prediction algorithms and used training data sets between tools, and phylogenetic differences in sequence diversity, physicochemical properties and evolved biological functions of AMPs between taxa, notable discrepancies may exist in performance between the currently available prediction tools. Here we tested if there is taxonomic bias in the prediction power in 10 tools with a total of 20 prediction algorithms in 19 invertebrate taxa, using a data set containing 1525 AMP and 3050 non-AMP sequences. We found that most of the tools exhibited considerable variation in performance between tested invertebrate groups. Based on the per-taxa performances and on the variation in performances across taxa we provide guidance in choosing the best-performing prediction tool for all assessed taxa, by listing the highest scoring tool for each of them.

## Introduction

Antimicrobial peptides (AMPs) are low molecular weight components of the innate immunity present virtually in all living organisms. Their importance has long been recognized, especially in the search for efficient therapeutic agents against multi-resistant bacteria ^1–3^. In the last few decades, invertebrates (mainly arthropods) have become central model organisms to identify and classify novel AMPs with potential utility in medicine ^4,5^. With the rapid accumulation of available genomic and proteomic data from a wide range of taxa and novel methods for the prediction of structural, physicochemical, and biological properties of peptides, the search for therapeutic-potential AMPs is one of the most exciting and promising areas in the life sciences ^6,7^ A In fact, by today a considerable number of AMP activity prediction tools has been built, mainly utilizing modern machine-learning algorithms to identify amino acid sequences with potential antimicrobial capacities, generally on the basis of their physicochemical parameters ^6,8^. Most often these parameters are some combinations of net charge, hydrophobicity, amphipaticity, isoelectric point, propensity to form certain structures (α-helix, ß-sheet, loops), polarity, and hydrophobic moment ^9,10^ and sometimes quantification of amino acid composition itself (e.g. as standard, normalized or pseudo composition, see ^11^ and references therein), which properties are associated with the capacity to attach to pathogen-specific targets (e.g. permeating bacterial cell membranes).

Beside to other challenges of AMP prediction using peptides from databases (see ^12^ and references therein), neglected taxonomic (and, indeed, phylogenetic) information may also contribute to biases in the training of prediction algorithms. Although AMP prediction tools are quite available, and some articles made attempts of benchmarking how efficiently these tools can identify AMPs in general (e.g. ^13^), it still remains overlooked whether or not these prediction tools may provide the same performance across the taxa from which putative AMPs are extracted. This could be rather problematic, owing to that even among phylogenetically related AMPs substantial differences may exist in the evolutionary trajectories of the amino acids between species. For example this might be expected to lead to differences in taxon-specific gene clustering (i.e. sequence diversity, affecting scalar physicochemical properties), or in the evolved functionality of peptides belonging to the given AMP family ^14–17^. Indeed, the importance of evolutionary origin of sequence diversity in these molecules was emphasized recently in invertebrates ^18^. While there is little knowledge about how such diversity may be associated with physicochemical properties and structural features of AMPs, it is not unreasonable to assume that unaccounted taxon-specific variation in them might affect the performance of prediction tools in recognizing antimicrobial function (e.g. due to taxon-based differences in the evolved physicochemical properties). Also, alternative tools using different methodological approaches and training sets further complicate the picture and may lead to discrepant performances, i.e. to differences between the prediction tools in how efficiently they recognize AMPs and non-AMPs in different taxa, which might depend on the taxonomic composition of the training data sets. In other words, the above mentioned factors may play a role in a low generalization-efficiency of the prediction algorithms that were trained on taxonomically (phylogenetically) limited training data set. Consequently, neglecting this aspect may lead to an increased incidence of both type I (falsely labelling non-antimicrobial peptides as (putative) AMPs) and type II (not recognizing true AMPs to have antimicrobial activity) errors when using inadequate tools.

This issue is anything but trivial, as a wide range of researchers rely on these AMP prediction tools and their outputs, from the search for invertebrate AMPs with therapeutical potential ^4^, to ecophysiological and ecoimmunological studies ^19^, and to the studies on the evolutionary history of invertebrate AMPs ^18^. If “taxonomic bias” exists in the prediction performance of available tools (i.e. if there are differences between prediction tools in how reliably they can distinguish between AMPs and non-AMPs in different taxa), a practical resolution (in terms of the usage of available prediction tools) could be the identification of which tool performs best for specific taxa, and consistently using these tools in the given groups. As a first approach to address this problem, taxon-specific assessment of the performance of currently available AMP prediction tools may help us to shed light on whether or not a taxon-bias is present at all. Accordingly, in the present study our aim is not to provide a comprehensive list of AMP prediction tools, or to introduce their algorithms and/or modern methods for AMP prediction, but to provide guidance in choosing the bestperforming prediction tool for specific taxa if taxonomic bias exists. In other words: our goal here is not to present the methodology and specific composition of training data sets in detail, but to assess whether currently available tools show considerable prediction bias between different invertebrate taxa, i.e. we don’t yet offer a specific solution to this technical problem at the level of algorithms, but highlight a yet neglected aspect of AMP prediction practice.

To do so, we acquired 1525 unique AMP sequences (from the protein database of the National Center for Biotechnology Information ^20,21^ and the Antimicrobial Peptide Database APD3 ^22^) and 3050 non-AMP sequences (from UniProt ^23^), from 19 invertebrate taxa, and used 10 freely available AMP prediction tools (with a total of 20 prediction algorithms), not only to assess the prediction tools’ overall performance, but their performance per taxa.

## Results

Overall ADAM showed the best performance with both of its algorithms (hidden Markov model, and support vector machine). Also, the artificial neural network classifier of CAMP_R3_ had relatively good performance, similarly to CS-AMPpred (all kernels, although the linear kernel did not perform as good as the polynomial and radial), and to iAMP-2L (Fig. 1). Based on F_1_ the weakest overall performance was observed in the case of StM, although based on Matthew’s correlation coefficient (MCC) the non-artificial-neural-network algorithms of CAMP_R3_ (discriminant analysis, random forest, and support vector machine) showed weakest performances.

**Figure 1.**
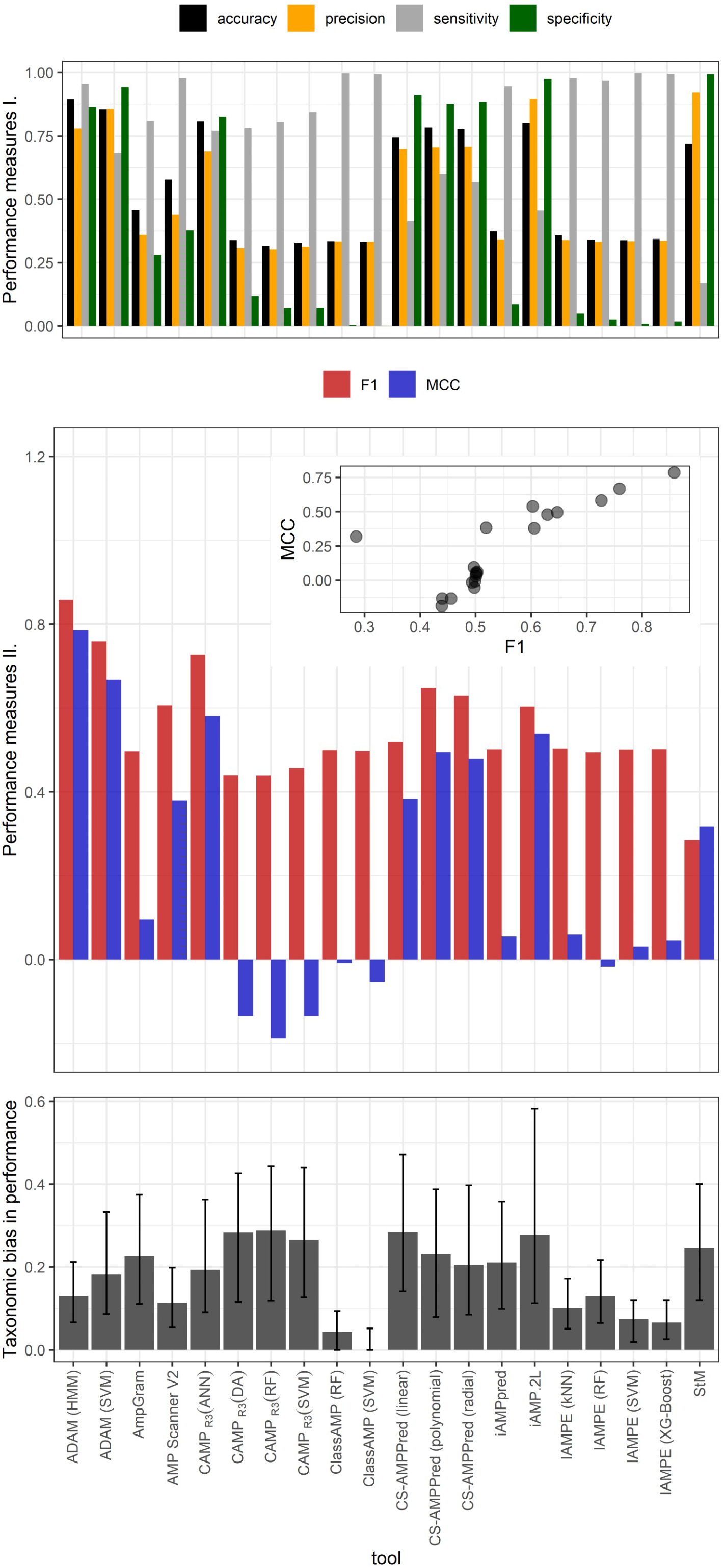
Classical performance measures (top panel), F_1_ scores and Matthew’s correlation coefficient (MCC; middle panel), and taxonomic bias in performance assessed as median of the distances in MCC scores between taxa, within a given prediction tool (bottom panel; see Methods for details); black segments on the bottom panel represent the value range between the first and third quartiles.

AMP prediction tools showed substantial variation in how well they performed in different taxa (Fig. 1 and 2). Class AMP and IAMPE classifiers showed almost no between-taxa variation, which was likely due to their poor performances, as they almost unanimously predicted AMP activity in all (even non-AMP) amino acid sequences. The largest taxonomic bias (i.e. largest median distance in tool performance between taxa) was observed in the random forest algorithm of CAMP_R3_ and in the linear kernel CS-AMPpred (Fig. 1); in these tools the discrepancy between the minimum and maximum value of MCC was 0.98, and 1.06, respectively. Even those prediction tools showing best performances based on their F_1_ and MCC scores had marked taxonomic biases: for example in the hidden Markov model ADAM the difference between minimum and maximum MCC was 0.44. Still, based on the F_1_ and MCC values calculated per taxa, this classifier yielded the highest performance for most of the taxa (Table 1).

**Figure 2.**
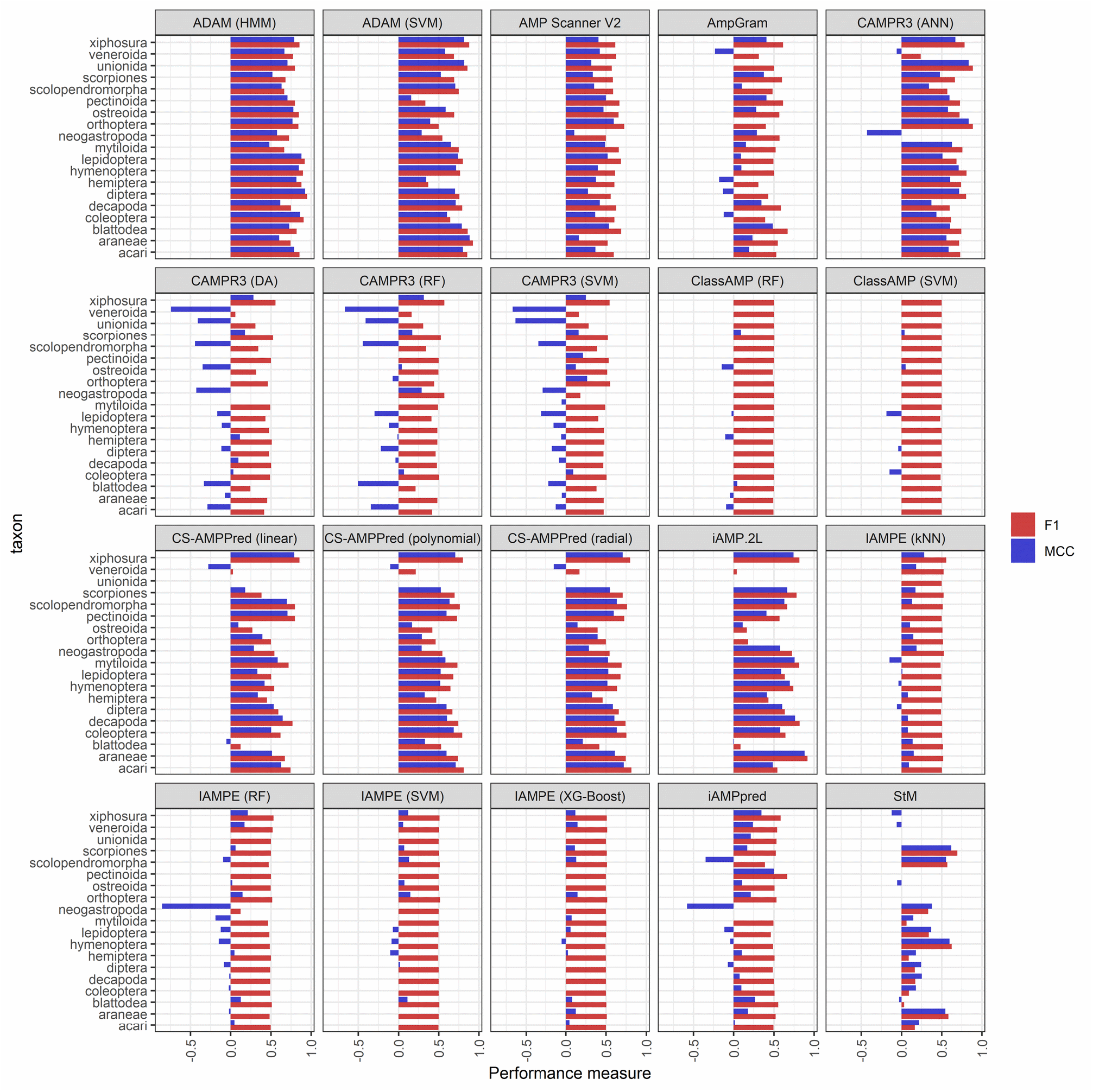
F_1_ and Matthew’s correlation coefficient (MCC) scores calculated for each taxon, separately within each prediction tool, in order to assess the prediction tools’ performance for each tested taxon.

**Table 1.** Best preforming prediction tools for given taxa based on F_1_ and Matthew’s correlation coefficient (MCC).

| Taxon | Best performing prediction tools |  |  |  |
| --- | --- | --- | --- | --- |
| | $F_1$ | | MCC | |
| Acari | ADAM (hidden Markov model) | 0.86 | ADAM (support vector machine) | 0.80 |
| Araneae | ADAM (support vector machine) | 0.93 | ADAM (support vector machine) | 0.89 |
| Blattodea | ADAM (support vector machine) | 0.86 | ADAM (support vector machine) | 0.79 |
| Coleoptera | ADAM (hidden Markov model) | 0.91 | ADAM (hidden Markov model) | 0.86 |
| Decapoda | iAMP-2L | 0.82 | iAMP-2L | 0.77 |
| Diptera | ADAM (hidden Markov model) | 0.95 | ADAM (hidden Markov model) | 0.93 |
| Hemiptera | ADAM (hidden Markov model) | 0.88 | ADAM (hidden Markov model) | 0.82 |
| Hymenoptera | ADAM (hidden Markov model) | 0.90 | ADAM (hidden Markov model) | 0.85 |
| Lepidoptera | ADAM (hidden Markov model) | 0.92 | ADAM (hidden Markov model) | 0.88 |
| Mytiloida | iAMP-2L | 0.82 | iAMP-2L | 0.76 |
| Neogastropoda | ADAM (hidden Markov model) | 0.73 | ADAM (hidden Markov model) | 0.58 |
| Orthoptera | CAMP<sub>R3</sub> (artificial neural network) | 0.89 | CAMP<sub>R3</sub> (artificial neural network) | 0.84 |
| Ostreoida | ADAM (hidden Markov model) | 0.85 | ADAM (hidden Markov model) | 0.78 |
| Pectinoida | ADAM (hidden Markov model) | 0.80 | ADAM (hidden Markov model) | 0.71 |
| Scolopendromorpha | CS-AMPpred (linear kernel) | 0.80 | ADAM (support vector machine) | 0.71 |
| Scorpiones | iAMP-2L | 0.78 | iAMP-2L | 0.67 |
| Unionida | CAMP<sub>R3</sub> (artificial neural network) | 0.89 | CAMP<sub>R3</sub> (artificial neural network) | 0.84 |
| Veneroida | ADAM (hidden Markov model) | 0.77 | ADAM (hidden Markov model) | 0.67 |
| Xiphosura | ADAM (support vector machine) | 0.88 | ADAM (support vector machine) | 0.82 |

In most cases taxonomic biases were tool specific, meaning that there were only a few taxa for which most of the prediction tools yielded consistently low or high performance (Fig. 3, and see also Supplementary Material Fig. S1). Specifically, we found negative bias for Veneroida, Unionida, and Neogastropoda (meaning that for these groups F_1_ and MCC scores were consistently low across prediction tools), and positive bias for Xiphosura and Araneae (meaning that for these groups F_1_ and MCC scores were consistently high across prediction tools). In taxa for which sample sizes were small (such as Unionida and Neogastropoda) this bias might simply be an artefact of chance, as any given prediction affects the estimated performance score much stronger than in taxa of larger sample sizes. However, in Veneroida and Araneae the sample sizes were not quite small, and in other small-sample-sized taxa no such pattern was observed. At this point it is difficult to find a satisfactory explanation for this observation. Given that most of the tested tools rely on amino acid sequence-based physicochemical measures, one possible cause might relate to the similarity of amino acid sequences in a given taxon to those used in the training sets of prediction tools. For example, even if a given taxon is not represented in a tool’s training set, similar physicochemical properties to the training sequences likely helps better recognition by the given tool. On the other hand, if physicochemical properties of a given taxon are significantly different from those taxa included in the training set, it will likely lead to poorer performance for that given query taxon not represented in the set. This should be addressed in future studies to clarify what mechanisms can lead to such biases in prediction performance.

**Figure 3.**
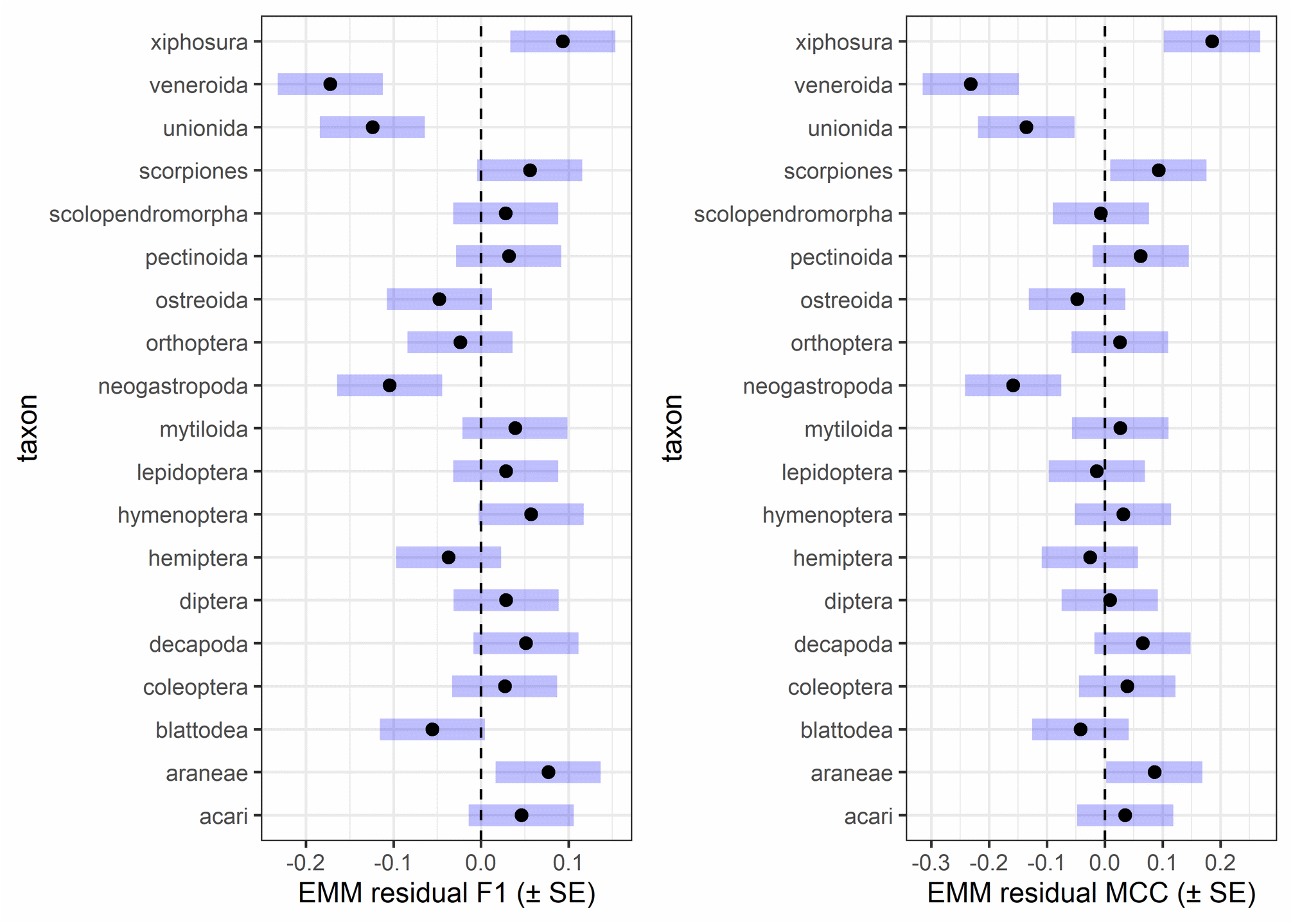
To check whether or not there were any taxa for which AMP predictions were consistently biased regardless of the used prediction tool we calculated “residual” F_1_ and Matthew’s correlation coefficient (MCC) scores. These residual measures are simply the difference between the F_1_ or MCC score of a given tool for a given taxon, and the arithmetic mean F_1_ or MCC for the given tool, averaged over all taxa within that tool. Hence, if for example a taxon would generally have low prediction successes across all tools, we would see that this taxon has consistently low (below zero) residual values in each tool. We would expect to see that these residual scores are normally distributed around zero when there is no such effect. To see if these residual F_1_ and MCC values have a mean significantly different from zero, estimated marginal means (EMM) were calculated by fitting a linear regression model with residual F_1_ and MCC as responses and taxon as categorical predictor. This figure depicts EMMs (black dots) and their standard errors (SE: blue horizontal bars); SE bars not crossing zero (dashed line) can be considered as the given EMM statistically significantly differing from zero (at α = 0.05).

## Discussion

The prediction tool ADAM using the hidden Markov model provided the best performance in most assessed taxa. While it is out of scope of this study to give in-depth insights on the algorithms behind the prediction tools, it should be noted that this tool’s strength likely comes from its set of homologues: it uses a probabilistic model based on which the already known AMPs are used to find the most likely homologue for the query peptide. This is a strength that few currently available prediction tools possess, namely that, instead of only using scalar physicochemical properties based on the amino acid sequence (which approach has its weaknesses, see Porto et al., 2020), sequences are classified based on their resemblance to known AMPs. Notably, such an approach may yield null-findings when presented a previously unknown AMP, but might potentially enable for accounting for evolutionary information. Although the hidden Markov model ADAM tool technically does not consider phylogeny for the taxa from which the query sequences originate, in future prediction tools homolog (and, indeed ortholog) finding may be a quite useful approach in the reliable *in silico* identification of AMPs. Surprisingly, tools accounting for (either species, or gene) evolution are still lacking, and studies addressing the question of functional typization of AMPs generally use amino acid composition (Porto et al., 2020), and/or quantitative structure-activity relationships ^24^. Whilst understanding the association between sequence, structure, and biological function is indisputably the forefront in the study of AMPs with therapeutical potential ^25,26^, evolutionary information may also help us in the identification (and, indeed, *de novo* design and synthesis) of candidate AMPs. Arguably, a taxonomically, or even phylogenetically informed perspective would be quite useful not only in the more robust and reliable identification of AMPs in invertebrates, but also in designing and engineering peptides with desirable properties (e.g. efficacy against antibiotic resistant bacteria, but low citotoxicity to eukaryotic cells), through a deeper understanding of how different antimicrobial properties evolved in AMPs of invertebrates. (For example, how evolutionary trajectories shaped AMPs to possess the advantageous physicochemical properties we observe today; see more on this concept in ^27^.) Hence, comparative phylogenetic analyses of AMPs would be an important aspect helping to understand the evolution of antimicrobial function in these molecules.

Substantial variation exists in how reliably given tools can predict AMP function of amino acid sequences belonging to different taxa. From a practical perspective this gives reason for caution when selecting the prediction tool for identifying putative AMPs from amino acid sequences of a given invertebrate taxon *in silico*. The substantial taxonomic bias in the different prediction tools’ performance may hint at important differences in the properties of amino acid sequences across taxa, but interpretation should be done with care, as this bias can have multiple origins. For example, various AMP families (e.g. defensins, cecropins) may have quite different physicochemical properties ^28^, and there may be substantial between-taxa variation in amino acid sequences of same AMP family ^18^. Although these aspects should be taken into account when working with AMPs, it is outside of the scope of our present study to go into details of the background and their possible resolutions.

We argue that our results can provide a firm basis for deciding which prediction tool(s) to use when working with invertebrate peptides. Joint consideration of tools’ performance and taxonomic bias may help to pick the best tools: for example, when one would like to test peptides of multiple taxonomical origins, it might be worth to consider tools with relatively low taxonomic bias and good performance, whereas when peptides originate from one known taxon, tool-specific performance in that group will be more important than the generalization capability of the given tool.

It should also be noted that our current approach is not without drawbacks. Firstly, the considerably uneven availability of AMPs for different taxa renders it difficult to consider our results robust for all the assessed taxa. As noted before, in taxa with small sample sizes (e.g. Neogastropoda and Unionida) F_1_ and MCC scores and prediction successes are more likely to be subjected to distortions by random chance than in taxa of larger sample sizes. Still, we think that as a practical guide our results are still of value even for taxa with small sample sizes, although with more caution. Secondly, one could propose that overlap between our test data set and training sets of prediction tools might also bias our performance estimates. However, we found no strong evidence for that, since neither the number nor the proportion of common amino acid sequences in our and the used tools’ data sets showed significant association with F_1_ scores, although the proportion of common sequences showed a weakly significant association with MCC (see Supplementary Material Fig. S2-S3). Notably, however, based on the supplementary figures, the moderate level of statistical significance (P = 0.029), and the results that in no other association did we find such connection renders it likely that this result is a false positive finding (type I error).

As previously mentioned, the different tools use different methodological approaches (e.g. CS-AMPpred uses support vector machine trained on 310 cysteine-stabilized peptides, utilizing scalar physicochemical properties, whereas ADAM hidden Markov model is based on profile models of more than 7000 AMPs and 759 identified structures; see Table 2 for additional details on the tested tools). Differences in the focus of AMP types undoubtedly restricts the potential range of peptides that can be recognized as to have antimicrobial function by a given prediction tool. It is also worth mentioning that taxonomic composition of the AMP and non-AMP sequence data sets used to train the different models might also shape the models’ performance, in terms of whether or not a tested putative AMP sequence comes from a taxon included in the training set. Unexpectedly, there was a weak trend that when a query taxon was present in the training set of a tool, then F_1_ and MCC scores tended to be slightly smaller (see Supplementary Material Fig. S4). In general, however, there was no significant association of the number and evenness of taxonomic groups in the training data sets with the tools’F_1_ and MCC scores (Supplementary Material Fig. S5).

**Table 2.**
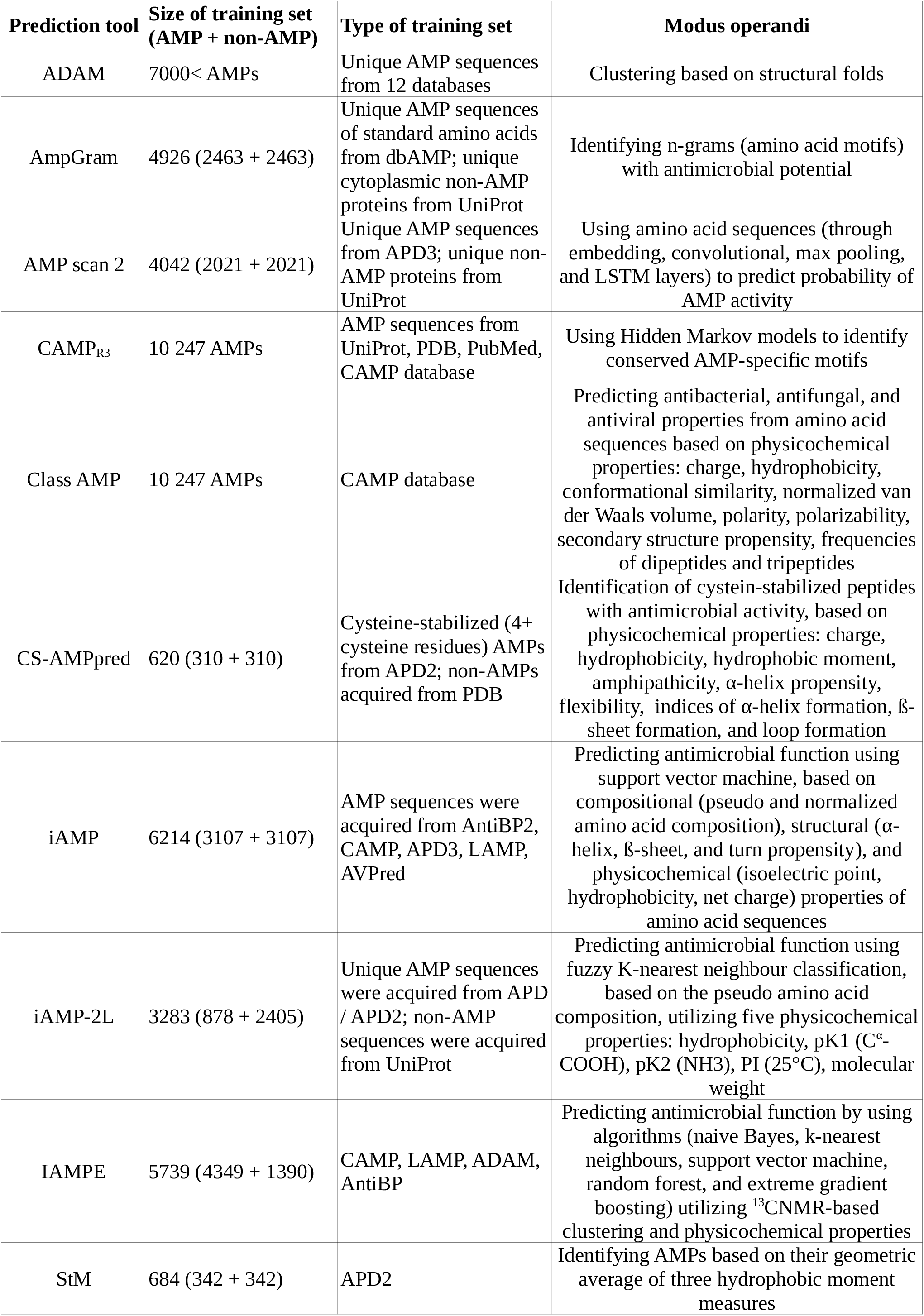
Additional informations about the used prediction tools. In the “Size of training set” column the number of sequences separately for AMPs and non-AMPs were not always available. In the “Type of training set” column where not indicated separately, both AMPs and non-AMPs were drawn from the name source(s).

| Prediction tool | Size of training set (AMP + non-AMP) | Type of training set | Modus operandi |
| --- | --- | --- | --- |
| ADAM | 7000< AMPs | Unique AMP sequences from 12 databases | Clustering based on structural folds |
| AmpGram | 4926 (2463 + 2463) | Unique AMP sequences of standard amino acids from dbAMP; unique cytoplasmic non-AMP proteins from UniProt | Identifying n-grams (amino acid motifs) with antimicrobial potential |
| AMP scan 2 | 4042 (2021 + 2021) | Unique AMP sequences from APD3; unique non-AMP proteins from UniProt | Using amino acid sequences (through embedding, convolutional, max pooling, and LSTM layers) to predict probability of AMP activity |
| CAMP<sub>R3</sub> | 10 247 AMPs | AMP sequences from UniProt, PDB, PubMed, CAMP database | Using Hidden Markov models to identify conserved AMP-specific motifs |
| Class AMP | 10 247 AMPs | CAMP database | Predicting antibacterial, antifungal, and antiviral properties from amino acid sequences based on physicochemical properties: charge, hydrophobicity, conformational similarity, normalized van der Waals volume, polarity, polarizability, secondary structure propensity, frequencies of dipeptides and tripeptides |
| CS-AMPpred | 620 (310 + 310) | Cysteine-stabilized (4+ cysteine residues) AMPs from APD2; non-AMPs acquired from PDB | Identification of cysteine-stabilized peptides with antimicrobial activity, based on physicochemical properties: charge, hydrophobicity, hydrophobic moment, amphipathicity, α-helix propensity, flexibility, indices of α-helix formation, β-sheet formation, and loop formation |
| iAMP | 6214 (3107 + 3107) | AMP sequences were acquired from AntiBP2, CAMP, APD3, LAMP, AVPred | Predicting antimicrobial function using support vector machine, based on compositional (pseudo and normalized amino acid composition), structural (α-helix, β-sheet, and turn propensity), and physicochemical (isoelectric point, hydrophobicity, net charge) properties of amino acid sequences |
| iAMP-2L | 3283 (878 + 2405) | Unique AMP sequences were acquired from APD / APD2; non-AMP sequences were acquired from UniProt | Predicting antimicrobial function using fuzzy K-nearest neighbour classification, based on the pseudo amino acid composition, utilizing five physicochemical properties: hydrophobicity, pK1 (C<sup>α</sup>-COOH), pK2 (NH<sub>3</sub>), PI (25°C), molecular weight |
| IAMPE | 5739 (4349 + 1390) | CAMP, LAMP, ADAM, AntiBP | Predicting antimicrobial function by using algorithms (naive Bayes, k-nearest neighbours, support vector machine, random forest, and extreme gradient boosting) utilizing <sup>13</sup>CNMR-based clustering and physicochemical properties |
| StM | 684 (342 + 342) | APD2 | Identifying AMPs based on their geometric average of three hydrophobic moment measures |

Additionally, due to the variable availability of AMP sequences for the different taxa, AMP family could not have been included when testing performance, even though there might be tool-specific differences in it for the different AMP families (i.e. a given tool may differ in its capacity to recognize defensins *versus* cecropins as AMPs). Notably, we know about no studies explicitly assessing the effect of AMP family on the efficiency of AMP prediction, therefore this aspect might be a quite relevant and important one to address in future studies. Furthermore, owing to practical reasons, non-AMPs (used both for training prediction tools, or for the assessment of their performance) are not standardized, i.e. these peptides are generally drawn randomly from those peptides that are available for the given taxa, with no known functions in antimicrobial immunity. Indeed, both in training and benchmarking data sets, false negative samples may be present in the form of yet-unknown (or simply undocumented) antimicrobial activity of some proteins used in the negative (non-antimicrobial) data sets. Also, non-overlapping biological functions between peptides from different taxa might introduce additional bias in the performance of prediction tools. For example, some membrane-active peptides might be expected to be falsely recognized as AMPs more often than others, due to their structural and physicochemical properties similar to that of AMPs ^6^. Hence, if the randomly drawn non-AMPs in a given taxon contain a higher proportion of such membrane peptides than in other taxa, this might confound the per-taxa-performance of the used prediction tool. Using standardized sets of non-AMP protein families (preferably of several, distinct biological functions) that have available sequence data for a wide range of taxa might help improve the performance of new prediction tools.

Although our current study addresses *in silico* prediction of antimicrobial activity in (invertebrate) peptides based on amino acid sequence, some of the details we have discussed are connected to the recent aspiration to improve upon currently used methodological standards of (pre)clinical testing of AMPs (such as in antimicrobial susceptibility testing, e.g. see ^29,30^). Indeed, more reliable and robust *in silico* prediction tools utilizing modern algorithms and ever larger data sets undoubtedly require standardization of quality of data. As such, AMP records could contain biologically and clinically relevant informations about the given peptides (as they have started to do so, see e.g. APD3 ^22^ or DRAMP ^31^). Similarly, standardized sets of non-AMPs characterized by laboratory-confirmed absence of antimicrobial activity in relevant contexts (*in vitro, ex vivo, in vivo*), and/or model organisms would quite likely help us improve current and future prediction tools and AMP design.

Overall, the results presented here highlight the necessity of taking taxonomic information into account when predicting antimicrobial function of putative invertebrate AMPs. At this point, although arguably crucial, finding the specific background of taxonomic bias (e.g. composition of training sets, or characteristics of prediction algorithms) in prediction performance of the assessed tools is outside the scope of our paper, and we are not able to offer technical details or solutions. We merely aimed to highlight an overlooked phenomenon, and to provide a practical guide to use the currently available and most popular tools. We hope that our results will serve as a useful guide in choosing the appropriate AMP prediction tool for testing candidate peptides in invertebrates to a wide range of researchers. This could be useful for the big-data / bioinformatic data style search for candidates in open databases and repositories, and for the identification of AMPs in invertebrates for ecophysiological studies as well. In the search for putative AMPs based on genomic and/or proteomic data, and immune profiling based on transcriptomics (i.e. screening produced peptides using transcriptome) in invertebrates may also benefit from our proposed approach. In addition, future studies to implement refined AMP prediction techniques, inclusion of taxonomic and/or phylogenetic information, utilization of explicit categorization of AMPs, and standardized sets of non-AMP sequences will undoubtedly help us to achieve more robust prediction of antimicrobial function in invertebrate models throughout our endeavour to identify peptides of potential therapeutical utility.

## Methods

All data handling and analyses were performed in the R software for statistical computing (ver. 4.0.1) ^32^. AMP sequences were acquired from the protein database of the National Center for Biotechnology Information ^20,21^ and the Antimicrobial Peptide Database APD3 ^22^ (specific keywords used in the search are listed in the Supplementary Material). Taxa for which we acquired AMP sequences were Acari (n = 76), Araneae (n = 45), Blattodea (n = 162), Coleoptera (n = 62), Decapoda (n = 64), Diptera (n = 447), Hemiptera (n = 63), Hymenoptera (n = 183), Lepidoptera (n = 136), Mytiloida (n = 31), Neogastropoda (n = 5), Orthoptera (n = 8), Ostreoida (n = 62), Pectinoida (n = 4), Scolopendromorpha (n = 10), Scorpiones (n = 103), Unionida (n = 4), Veneroida (n = 48), and Xiphosura (n = 12). The quantiles for length of amino acid sequences were: Q_min_ = 8, Q_25_ = 39, median (Q_50_) = 63, Q_75_ = 95, Q_max_ = 452.

Non-AMP sequences were downloaded from UniProt ^23^: for each taxon, a large batch of proteins without known antimicrobial activity was acquired, between the length of 10 and 200 amino acids (for specific search expression see Supplementary Material), from which 2n for each taxon were randomly drawn (where “n” refers to the number of acquired AMP sequences in the given taxa), without duplicate sequences. Amino acid sequences and data tables used in our study have been made available at figshare (https://figshare.com/projects/Taxonomic_bias_in_AMP_prediction/114585).

After the acquisition, both AMPs and non-AMPs were used with 10 AMP prediction tools; as some of the tools included multiple algorithms, in total there were 20 algorithms for the prediction of AMP properties for each acquired sequence (for the detailed list of the used prediction tools see Table 3).

**Table 3.**
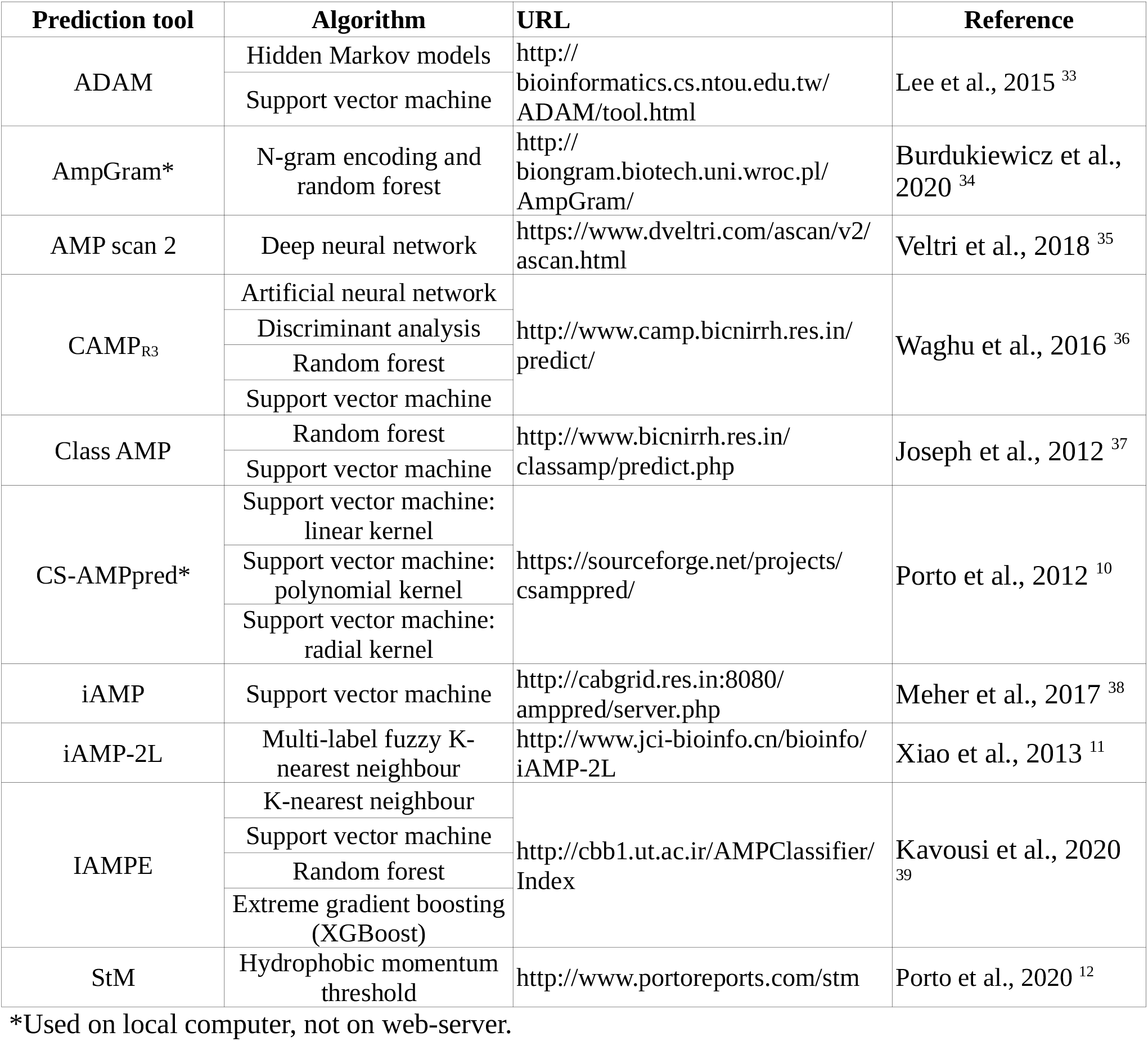
List of the used prediction tools (and algorithms within some tools).

In the case of prediction tools providing a probability for a given amino acid sequence to be an AMP, we used the conventional 0.5 threshold (i.e. sequences with >50% probability were considered to be classified as AMP). Results from the prediction tools were evaluated using the typical binary classification performance measures:

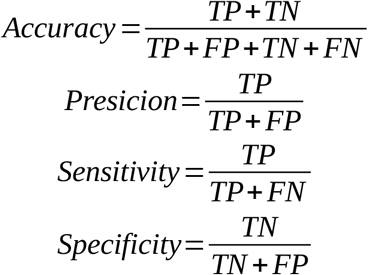

where TP, TN, FP, and FN stand for True Positive, True Negative, False Positive, and False Negative prediction, respectively. We also calculated the F_1_ score, a standard measure of classification efficiency, as the harmonic mean of precision and sensitivity ^40^:

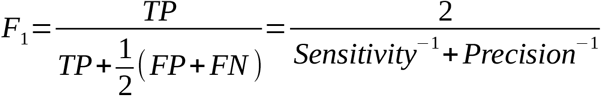

F_1_ scores can range between 0 and 1, the former indicating that either sensitivity and/or precision is zero, whereas the latter indicates perfect precision and sensitivity. Furthermore, we have calculated Matthew’s correlation coefficient (MCC) as well, using the R-package “mltools”^41^. MCC was found to be unbiased in imbalanced confusion matrices in binary classification, and was suggested to be adopted as the standard performance measure in the evaluation of binary classification tasks ^42^. MCC values range between −1 and 1: larger positive values reflect better prediction, zero indicates prediction no better than random chance, and negative values indicate consistent disagreement between prediction and observation. In our evaluations and analyses we decided to report both F_1_ scores and MCC, because F_1_ is still widely used and a large number of researchers may find it helpful to see both performance measures while interpreting our findings.

Tool performances were also assessed separately for taxa. When inferring on the best performing tool for each taxon, we used the “highest score” rule of thumb, and we listed the prediction tools both with the highest F_1_ score and highest MCC for each specific taxon. We also presented all scores for all tool-taxon combinations (see Results). Additionally, taxonomic bias in prediction power of tools was calculated as the median distance between MCC scores within taxa, separately for each prediction tool. The utilized measure of taxonomic bias is expected to range between 0 and 2, with smaller values indicating no or small differences in MCC across taxa within a given prediction tool, whereas larger values indicate substantial variation in MCC across taxa within the given prediction tool.

## Acknowledgements

We are indebted for the constructive comments and suggestions of two anonymous reviewers. The work was partly supported by the project for development of Complex Health Multidisciplinary Competence Center, Debrecen, Hungary (GINOP-2.3.4-15-2020-00008).

## Author contributions

R.Z. did the conceptualization and design of the study, and the main analyses. All authors were involved in data collection and handling. R.Z. drafted and completed the manuscript. All authors were involved in reviewing and finishing the review.

## Notes

https://figshare.com/projects/Taxonomic_bias_in_AMP_prediction/114585

### Competing Interest Statement

The authors have declared no competing interest.

