## Supplementary Material for "Taxonomic bias in AMP prediction of invertebrate peptides"

#### *Antimicrobial peptides*

In APD3 acquisition was simply: typing "invertebrate" in the "Name" box, then copying the result table.

Search expressions used at **NCBI** protein database:

- ((defensin) AND flies[porgn:__txid7147] NOT like NOT putative NOT predicted NOT precursor NOT partial)
- ((cecropin) AND flies[porgn:__txid7147] NOT like NOT putative NOT predicted NOT precursor NOT partial)
- ((diptericin) AND flies[porgn:__txid7147] NOT like NOT putative NOT predicted NOT precursor NOT partial)
- ((attacin) AND flies[porgn:__txid7147] NOT like NOT putative NOT predicted NOT precursor NOT partial)
- ((drosocin) AND flies[porgn:__txid7147] NOT like NOT putative NOT predicted NOT precursor NOT partial)
- ((defensin) AND hymenopterans[porgn:__txid7399] NOT like NOT putative NOT predicted NOT precursor NOT partial)
- ((diptericin) AND hymenopterans[porgn:__txid7399] NOT like NOT putative NOT predicted NOT precursor NOT partial)
- ((defensin) AND bugs[porgn:__txid33345] NOT like NOT putative NOT predicted NOT precursor NOT partial)
- ((diptericin) AND bugs[porgn:__txid33345] NOT like NOT putative NOT predicted NOT precursor NOT partial)
- ((defensin) AND moths[porgn:__txid7088] NOT like NOT putative NOT predicted NOT precursor NOT partial)
- ((cecropin) AND moths[porgn:__txid7088] NOT like NOT putative NOT predicted NOT precursor NOT partial)
- ((cecropin) AND beetles[porgn:__txid7041] NOT like NOT putative NOT predicted NOT precursor NOT partial)
- ((attacin) AND beetles[porgn:__txid7041] NOT like NOT putative NOT predicted NOT precursor NOT partial)

- ((holotricin) AND beetles[porgn:__txid7041] NOT like NOT putative NOT predicted NOT precursor NOT partial)
- ((coleoptericin) AND beetles[porgn:__txid7041] NOT like NOT putative NOT predicted NOT precursor NOT partial)
- ((coprisin) AND beetles[porgn:__txid7041] NOT like NOT putative NOT predicted NOT precursor NOT partial)
- ((defensin) AND roaches[porgn:__txid85823] NOT like NOT putative NOT predicted NOT precursor NOT partial)
- ((defensin) AND mites & ticks[porgn:__txid6933] NOT like NOT putative NOT predicted NOT precursor NOT partial )
- ((defensin) AND spiders[porgn:__txid6893] NOT like NOT putative NOT predicted NOT precursor NOT partial )
- ((defensin) AND scorpions[porgn:__txid6855] NOT like NOT putative NOT predicted NOT precursor NOT partial )
- ((defensin) AND molluscs[porgn:__txid6447] NOT like NOT putative NOT predicted NOT precursor NOT partial )
- ((termicin) AND roaches[porgn:__txid85823] NOT like NOT putative NOT predicted NOT precursor NOT partial )
- ((hemipterícin) AND bugs[porgn:__txid33345] NOT like NOT putative NOT predicted NOT precursor NOT partial )
- ((defensin) AND dragonflies & damselflies[porgn:__txid6961] NOT like NOT putative NOT predicted NOT precursor NOT partial )
- ((defensin) AND grasshoppers, crickets, and katydids[porgn:__txid6993] NOT like NOT putative NOT predicted NOT precursor NOT partial )
- ((diptericin) AND grasshoppers, crickets, and katydids[porgn:__txid6993] NOT like NOT putative NOT predicted NOT precursor NOT partial )
- ((attacin) AND grasshoppers, crickets, and katydids[porgn:__txid6993] NOT like NOT putative NOT predicted NOT precursor NOT partial )
- ((phormicin) AND flies[porgn:__txid7147] NOT like NOT putative NOT predicted NOT precursor NOT partial)

***Non-antimicrobial peptides***

Search expressions used at UniProt protein database was based on the template:

taxonomy:TAXON length:[10 TO 200] NOT partial NOT putative NOT predicted NOT precursor NOT hypothetical NOT keyword:antimicrobial

where “TAXON” was replaced with one of the taxon names below, separately for each taxon:

- acari
- araneae
- blattodea
- coleoptera
- decapoda
- diptera
- hemiptera
- heteroptera
- hymenoptera
- lepidoptera
- mollusca
- mytiloida
- neogastropoda
- orthoptera
- ostreoida
- scolopendromorpha
- scorpiones
- xiphosura

### Supplementary Figure S1

To check whether or not there were any taxa for which AMP predictions were consistently biased regardless of the used prediction tool we can calculate a sort of “residual” $F_1$ and $MCC$ scores. These measures are simply the difference between the $F_1$ or $MCC$ score of a given tool for a given taxon, and the arithmetic mean $F_1$ or $MCC$ for the given tool, averaged over all taxa within that tool. Hence, if for example a taxon would generally have low prediction successes across all tools, we would see that this taxon has consistently low (below zero) residual $F_1$ and $MCC$ values in each tool. However, we would expect to see that these residual scores are normally distributed around zero when there is no such effect.

The figures below depict density kernel distributions of residual $F_1$ and $MCC$ values separately for taxa; dashed vertical lines denote zero, and solid vertical lines denote arithmetic mean.

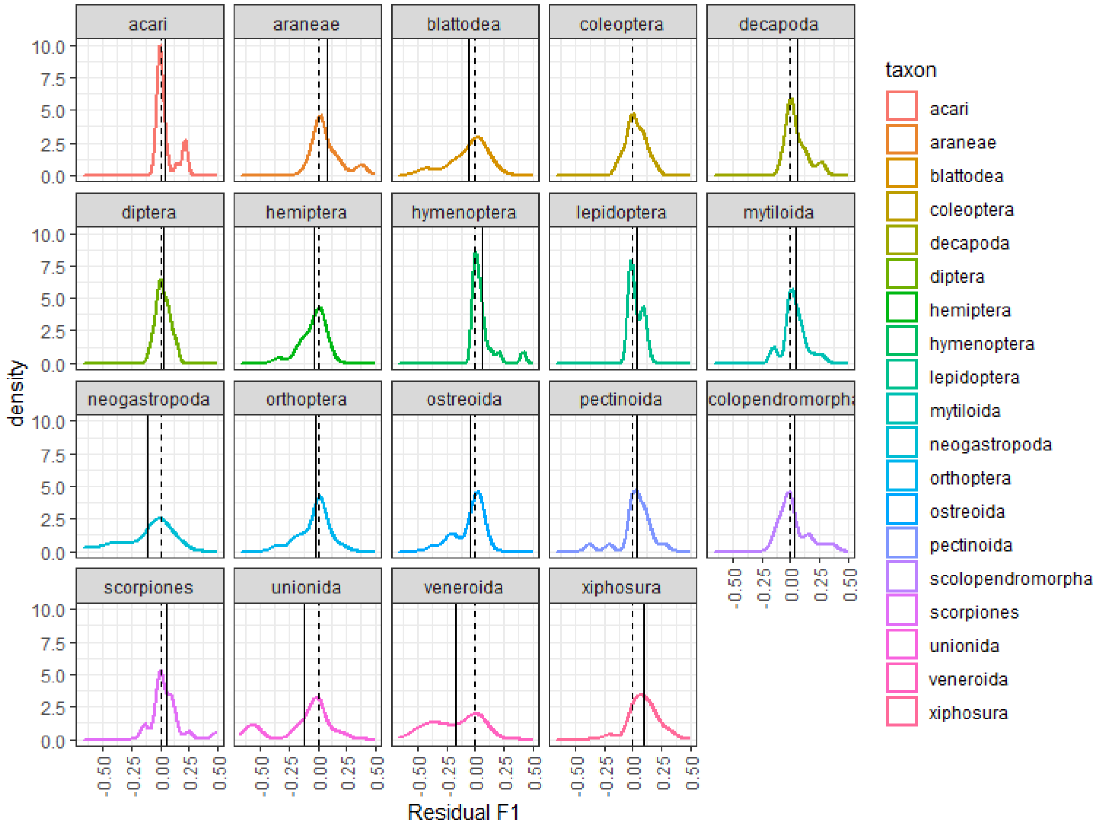

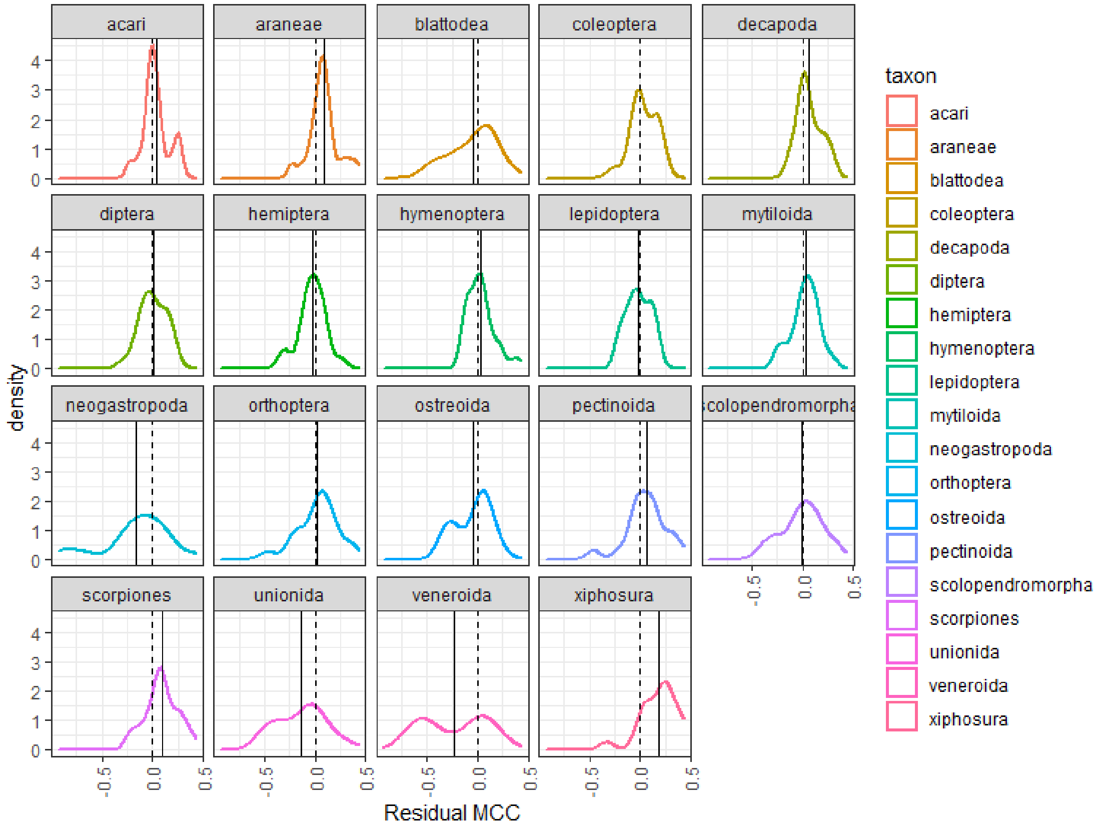


Note that this is different from assessing the so-called taxonomic bias in tools' prediction efficiency, because in this case we assess whether or not there are specific taxa for which most prediction tools give biased results (whereas in the taxonomic bias we assess prediction performance within, rather than across, tools).

### Supplementary Figure S2

Overlap in amino acid sequences (both AMP and non-AMP) between prediction tools' training sets and our test set. The proportion of common sequences in training set shows how big proportion of the training set amino acids were found in our test set.

To assess the taxonomical composition of the training data sets of the tested prediction tools we downloaded the whole training sets, and used BLAST+ (Camacho et al., 2009) to identify the taxonomical order of organism to which given amino acid sequences belonged. Admittedly, this work-around does not yield 100% matches, but it was still a relatively quick and reliable approach to acquire taxonomic data for sequences without such information.

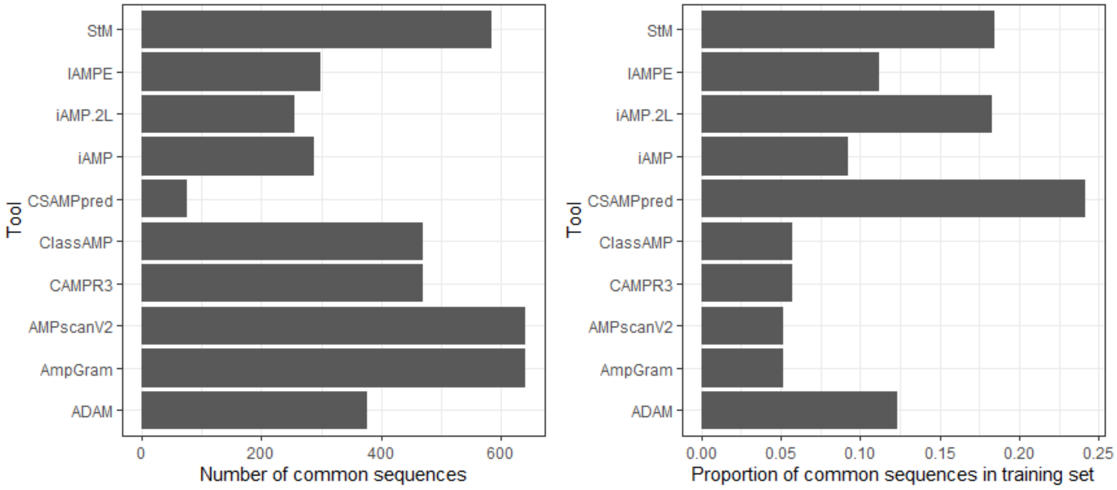

### Supplementary Figure S3

Association of performance (as $F_1$ and $MCC$ scores) with the number and proportion of amino acid sequences overlapping between our test set and the prediction tools' training sets. To test statistical significance of these associations, mixed-effects linear regression models were fitted, in which the response variables (in two separate models) were the prediction tool-specific $F_1$ and $MCC$ scores, and (in separate models) the predictors were the number and proportion of overlapping sequences; we defined the tools' training set pools as random effect. (The models were fitted with the R-package *glmmTMB* (Brooks et al., 2017).)

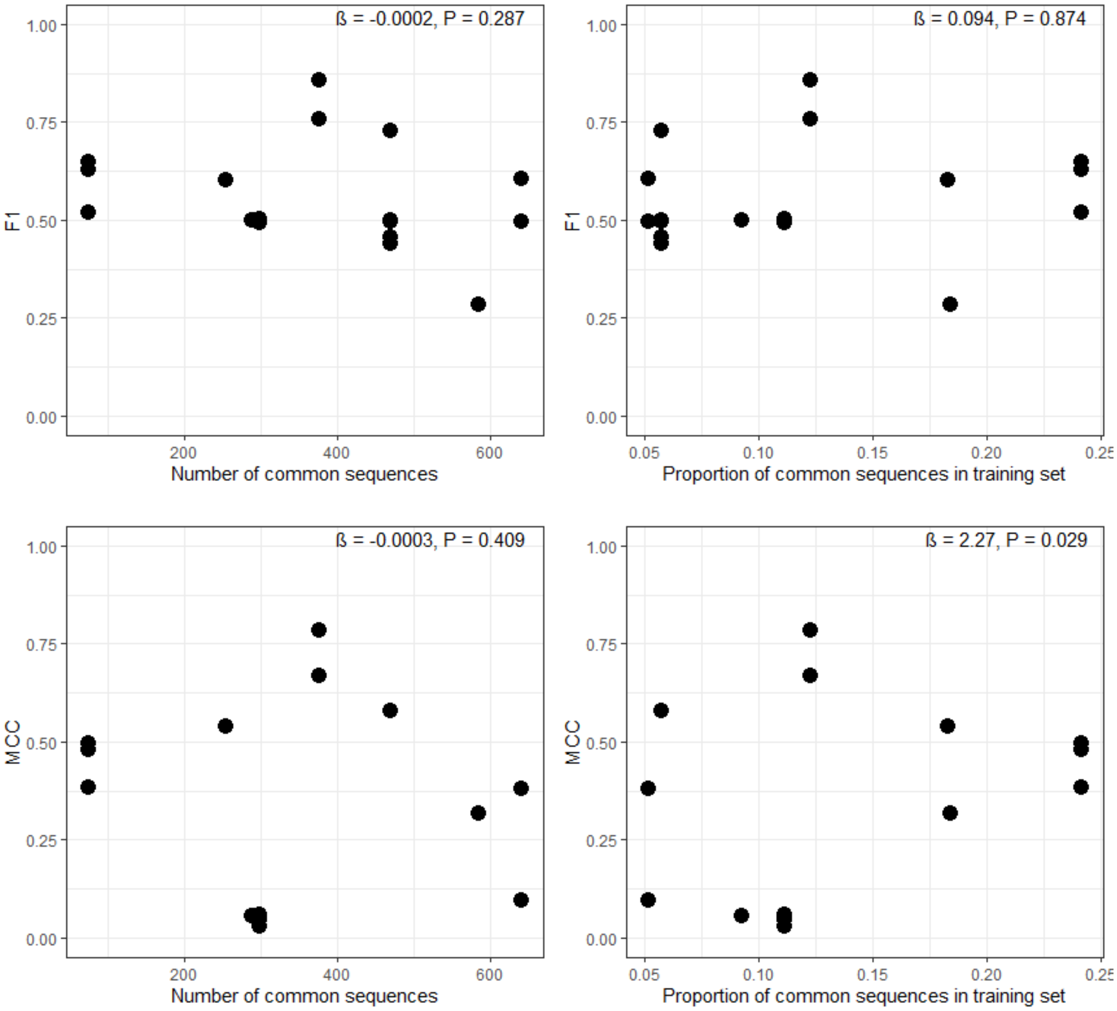

### Supplementary Figure S4

Checking whether the presence of the query taxon in the prediction tool's training set affects $F_1$ or $MCC$ scores. To test the statistical significance of this effect, a mixed-effects linear regression models were fitted (with *glmmTMB*), separately for $F_1$ and $MCC$ as response, query taxon presence in tools' training sets as predictor factor, and the tools' training set pools as random effect.

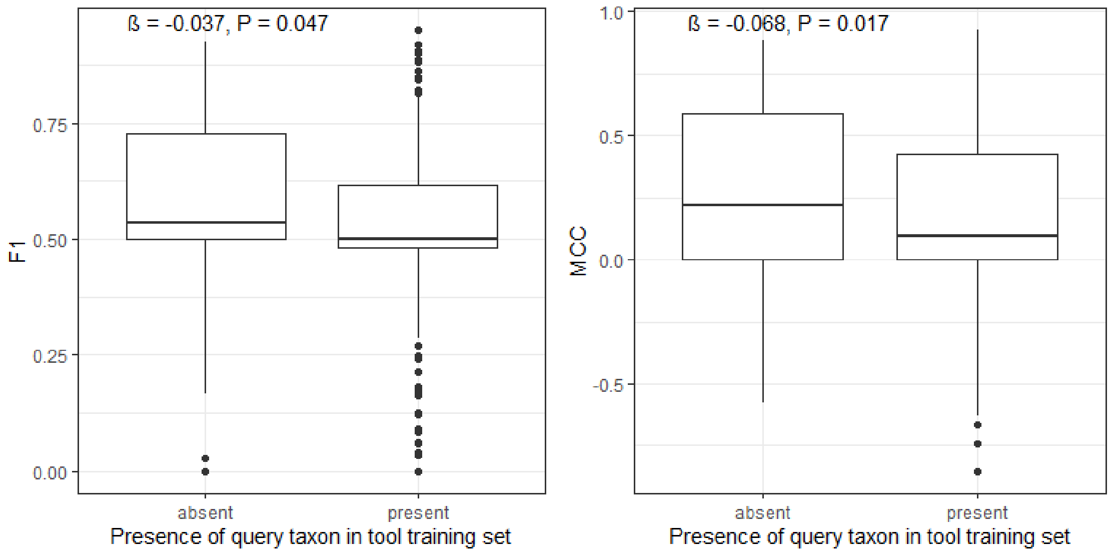

### Supplementary Figure S5

Associations of $F_1$ and $MCC$ with number and evenness of invertebrate taxonomic orders in the training data sets. Evenness is quantified as the normalized entropy (also known as "efficiency") of the proportion of taxa within a given tool:

$$E = \sum_{i=1}^{n} \frac{p(x_i) log(p(x_i))}{log(n)}$$

where **E** is evenness, **n** is the number of taxa in the given prediction tool's training set, **x** is the number of amino acid sequences in the training set belonging to the given taxon, and **p(x)** is the proportion of the given taxa in the data set.

To test the statistical significance of these associations, mixed-effects linear regression models were fitted (with *glmmTMB*), with $F_1$ and $MCC$ as responses in separate models, and (also in separate models) number and evenness of taxa (order) as predictors; the tools' training set pools was set as random effect.

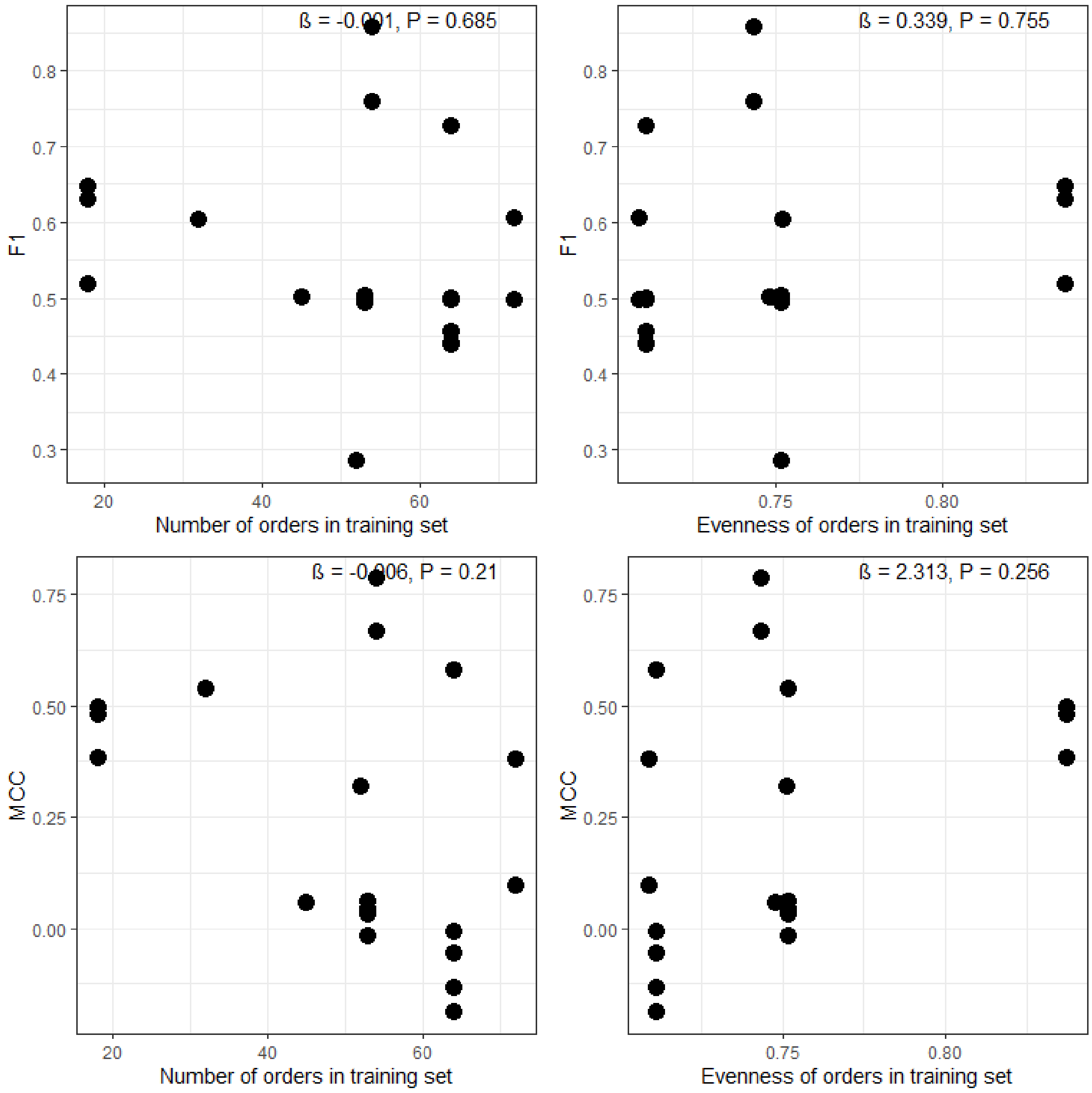
